# Effective Cell Immunoablation in Undisrupted Developing Avian Embryos

**DOI:** 10.1101/091116

**Authors:** Maríacruz López-Díaz, Julia Buján-Varela, Carlos Cadórniga-Valiño

## Abstract

In birds the construction of germline chimeras by grafting exogenous primordial germ cells (PGCs) during embryonic development is feasible since they migrate to the gonads through the blood. Up to date, the efficiencies are highly variable, in part dependent on the destruction of endogenous PGCs in the recipient embryo. We show an almost complete ablation of the endogenous PGCs in stage X embryos using a baby rabbit serum (BRS), with previous cellular signaling by specific antibodies (SSEA1). The application of the treatments, either on epiblast or subgerminaly, produced the reduction of the PGCs in the embryos in a dose dependent manner. No malformations or damages were detected in the treated embryos. However, subgerminal injection of this cocktail produced a massive cellular destruction in all embryos. Therefore, sequential application is a selective and effective method to produce receptor embryos. Nevertheless, it can also be highly destructive if the mixture is applied locally, this could be useful in the treatment of malignancies.

**SUMMARY STATEMENT:** An immunosurgery procedure is described that yields an almost complete ablation of primordial germ cells in early developing chick embryos, thus increasing the expected rates of chimerism when foreign PGCs are grafted onto these embryos

## INTRODUCTION

Very few studies have been done “in vivo” since 1975 when Solter and Knowles coined the term of “immunosurgery”. These authors were able to ablate the trophectoderm cells in mouse embryos by using antibody and complement. In 2007 Chen and Melton by using antibodies against red blood cells and heterologous guinea pig serum as a source of complement, achieved successfully isolation of the inner cell mass in human embryos. Also Gerhart et al. developed an efficient ablation procedure of two epiblast cell types with neuronal and muscular epitopes by using specific antibodies and baby rabbit serum (BRS) in stage X chicken embryos (Gerhart et al 2008, 2010).

On the other hand, in birds it is needed to develop new assisted reproduction procedures since avian oocytes and embryos are organized as an extremely precise biological structure, the egg. This complex assembly of oocyte/embryo does not stand freezing as is the case of mammalian embryos. Every aspect of the fertilized egg (pH of the different compartments, gas permeability, density relations and floatability, access to the different layers of nutrients, etc) is finely adjusted (Stern 1991, Lopez-Diaz et al. 2016). However, avian embryo has morpho-physiological peculiarities which allow the spread of a genotype or a specific lineage from the precursor cells of sperm and oocytes, the primordial germ cells (PGCs), which can be cryopreserved ex-situ. This assisted reproduction procedure involves the construction of germline chimeras by grafting, during embryonic period, PGCs of the desired lineage on a recipient embryo of a different strain (van de Lavoir 2006).

The inactivation of the recipient's germ line has been used as a strategy to improve the degree of germinal chimerism. For decades, the commonly used methods have been irradiation (Maeda et al, 1998, Carsience et al, 1993; Kino et al 1997; Speksnijder and Ivarie, 2000; Lia et al, 2001; Nakamura et al, 2012) or administration of cytotoxic agents (Hemsworth & Jackson 1963; Reynaud 1977; Mozdziak et al 2006; Song et al, 2005; Petitte et al 1990; Bresler at al., 1994; Aige Gil V and Simkiss K 1991, Naito et al. 2015). Both are effective, but completely unspecific, equally affecting all populations of embryonic cells, damaging the viability of the recipient embryo. In fact, the results usually achieved have an extremely high variability in the transmission rates of the grafted germline (2.9-100%), whether the endogenous PGCs are inactivated (Naito et al. 2015) or not (van de Lavoir 2006, White et al. 2015), this means that the procedure is not under control.

The origin and migration of avian PGCs have been well characterized (Dubois 1969; Kuwana and Fujimoto, 1984; Muniesa and Dominguez, 1990) since first described by Swift (1914). It has been estimated that the number of PGCs at oviposition (stage X embryo) is 100-120. Such a reduced number of cells and the high potential of developmental plasticity make this developmental stage the best time for the elimination of endogenous PGCs.

On the other hand, the combined cytolytic action of antibodies and complement has been well known for over a century (Wiemann 1994) and has been applied with different therapeutic and experimental purposes. In rabbits previously grafted with cancer cells (Kalfayan & Kidd, 1953) the combination of serum complement with antibodies against Brown-Pearce carcinoma cells achieved effective inhibition of tumor growth. Nowadays, one of the best treatments against cancer is the use of antibodies, with over 20 of different nature approved by various regulatory agencies (Macor & Tedesco 2007). As a consequence, in the last few decades the antibody-complement actions have been deeply investigated. It is known that complement modulates the complement dependent cytotoxicity (CDC) and antibody dependent cytotoxicity (ADCC). However, complement can have opposite effects; it can favor CDC and inhibit ADCC. Our understanding of the mechanisms involved when antibody and complement are acting together is still in its infancy (Rogers et al. 2014).

Like many malignant neoplastic cells, PGCs have a glycocalix rich in trisaccharide Galβ (1-4), Fucα (1–3)-GlcNAcβ1-R4 which is recognized by antiSSEA1. In fact, antiSSEA1 has been widely used to enrich selectively PGCs from stage X embryos (Etches 1998) and gonocytes from embryos of 5days, stage 27 (Mozdziak 2006).

Therefore, the objective of this study was to create viable chick embryos without endogenous PGCs, taking advantage of the complement dependent cytolysis (CDC). We used antiSSEA1, Ig M monoclonal antibody, together with baby rabbit serum (BRS) as a source of complement to induce a selective ablation of the PGCs in stage X chicken embryos. These chicken embryos could be used as receptors for unique exogenous PGCs.

## RESULTS

### “In vitro” cytolysis of avian blastoderm cells

*Competition tests of dispersed blastodermal cells with haemolytic system (HS) composed by sheep red blood cell (SRBCs) and haemolysin in gelatine-Veronal buffer with Ca*^*2+*^ *Mg*^*2+*^. Haemolytic complement activity measures the complement classical pathway in serum. Fifty percentage of haemolytic complement activity (CH_50_ or Kd=dissociation constant) in competition curves, measures serum haemolysis capacity in SRBCs. CH_50_ is sensitive to any reduction of the classical pathway components. Therefore, haemolysis absorbance of SBRCs was evaluated (y-axis) with decreasing concentrations of baby rabbit serum (BRS) as a complement source (x-axis), without antiSSEA1 (yellow line) and with antiSSEA1 1/100 (blue line) in presence of blastodermal cells. Saturation curve with dispersed blastodermal cells and antiSSEA1 1/100 revealed that, when blastodermal cells and antiSSEA1 were present, the saturation was reached and that the Kd (CH_50_) was between 1/10 and 1/100 dilutions of BRS (Fig 1a). These dilutions were used for selective PGCs ablation studies. We performed next experiments focused on dilutions of 1/00 and 1/40 for antiSSEA1 and BRS, respectively.

*Evaluation of cellular lysis with trypan blue after challenges with antiSSEA1 and BRS*. In isolated blastoderms the cellular lysis was evident in zona pellucida, where PGCs are localized at this stage, while in control embryos none of the cells were stained (Fig 1b, A and B).

**Figure 1.**
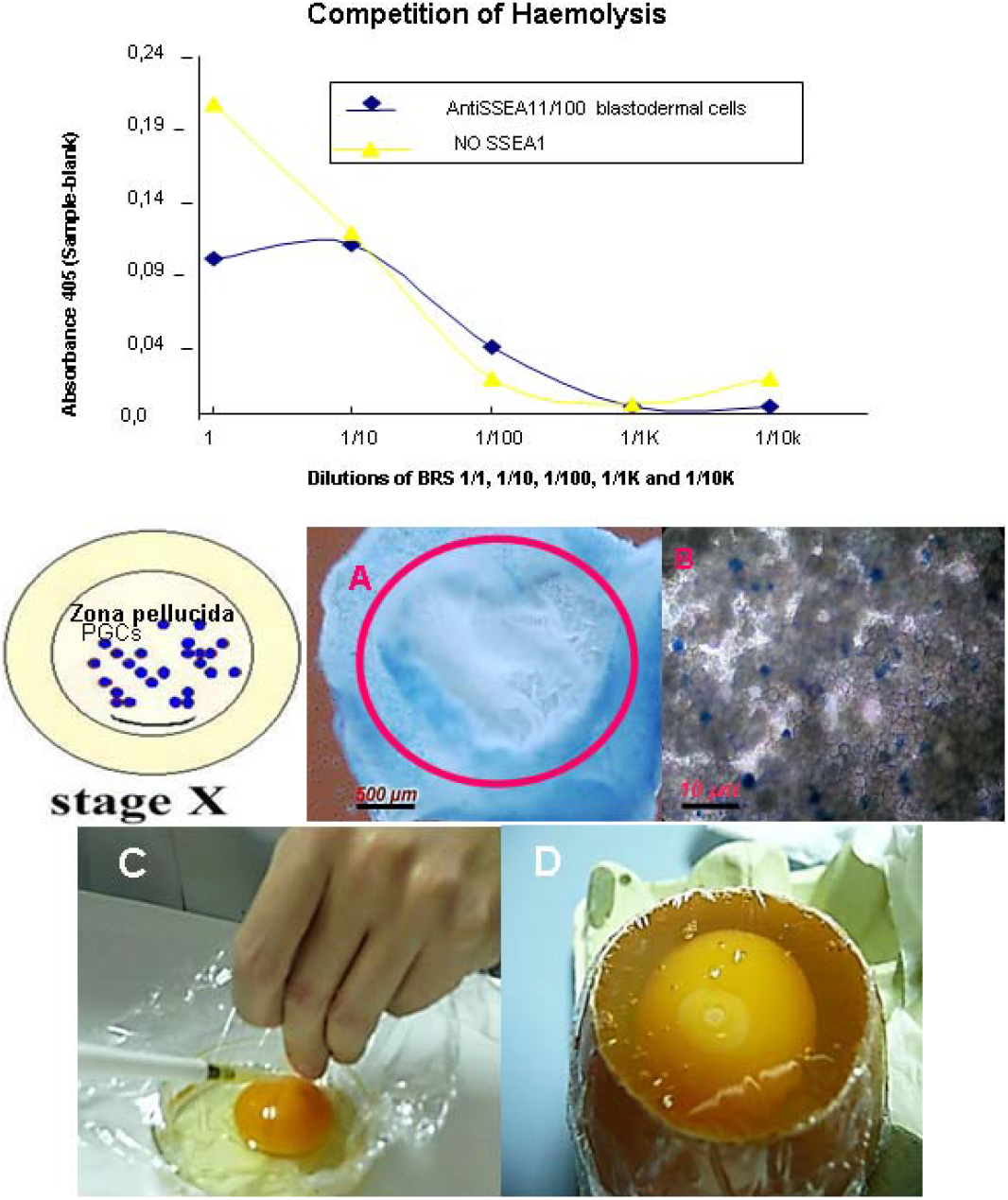
1a Saturation curve. Competition test with dispersed blastodermal cells and anti SSEA1 1/100. Haemolysis absorbance of sheep red blood cell (SBRC y-axis) with decreasing concentrations of baby rabbit serum (BRS) as a complement source (x-axis), without antiSSEA1 (yellow line) and with antiSSEA1 1/100 (blue line). When blastodermal cells and antiSSEA1 were present the saturation was reached and the CH50 (Kd) was between 1/10 and 1/100 dilutions of BRS. Blastoderm cell suspensions from ten stage X embryos were used in each saturation curve. **1b. Deposition of antiSSEA1 and baby rabbit serum to the dorsal side of epiblast and evaluation of cellular lysis with trypan blue**. Chicken embryos stage X (Ax4 and B x200), A) Control embryo treated with PBS and trypan blue exposed, none cell was stained by trypan blue (stereoscope Leica MZIII) B) Zona pellucida of one embryo challenged with antiSSEA1 and baby rabbit serum, there are blue cells stained (NIKON, Eclipse TE300, inverted microscope). C) Deposition of the treatment to the dorsal side of epiblast. D) 24 hours embryo, developmental stage referred in the manuscript as “early development”.

### Ex-ovo treatments applied to stage X embryos and incubated in surrogated egg shells following Perry’s system II

*Embryonic development and number of PGCs in embryos treated with deposition of antiSSEA1 and baby rabbit serum to the dorsal side of epiblast*. Two times of embryo development were evaluated, “early development” 24 hours after the application of the treatment (Fig 1b C and D) and “late development” 3-day. The evaluation of embryonic development revealed that the treatments and transfer manipulations did not affect neither embryo “early development” (67-100%) nor “late development” (56-90%) in any of the treated groups (AntiSSEA1 10/C+, 100/C+ and 1000/C+ groups) compared with control groups (AntiSSEA1-/C- and AntiSSEA1-/C+), Table 1. The mean survival days was 4.7, as expected in normal embryo development following Perry’s system II (Fig 2a), with no significant differences between treatments. However, the number of PGCs detected was decreased in treated (4, 2.7 and 4 PGCs in AntiSSEA1 10/C+, 100/C+ and 1000/C+ groups, respectively) compared with control embryos (50 and 45 PGCs in AntiSSEA1-/C- and AntiSSEA1-/C+) (Fig 2b). A total of 2,400 tissue sections were immunostained and evaluated.

**Table 1.**
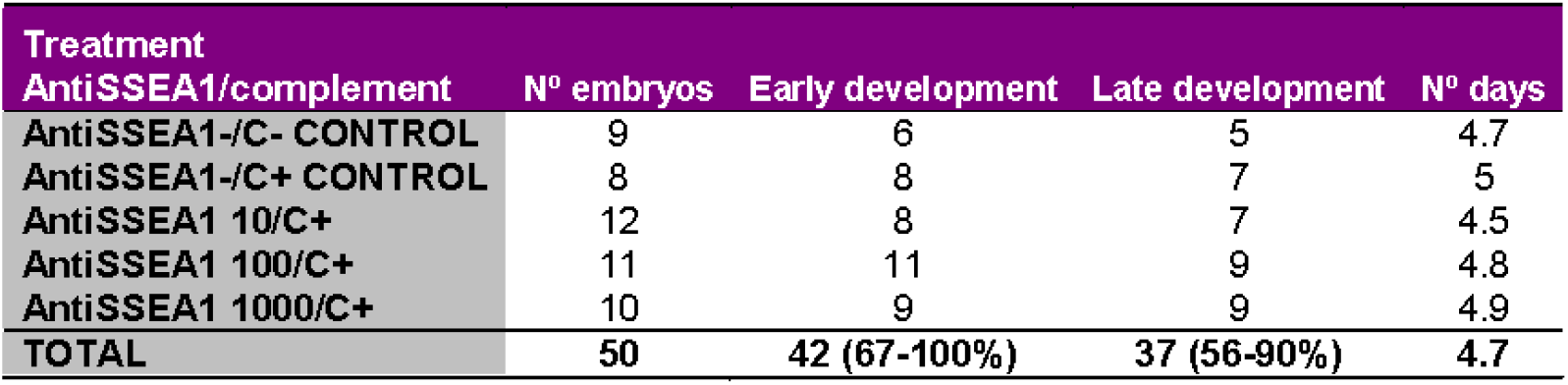
Embryos distribution of “early” and “late development” when the treatment was deposited to the dorsal side of epiblast at stage X. “Early development” 24 hours after the application of the treatment, the embryo continues to advance. “Late development” at day fourth embryos with heartbeat and vascular tree well formed.

| Treatment | Nº embryos | Early development | Late development | Nº days |
| --- | --- | --- | --- | --- |
| AntiSSEA1-/C- CONTROL | 9 | 6 | 5 | 4.7 |
| AntiSSEA1-/C+ CONTROL | 8 | 8 | 7 | 5 |
| AntiSSEA1 10/C+ | 12 | 8 | 7 | 4.5 |
| AntiSSEA1 100/C+ | 11 | 11 | 9 | 4.8 |
| AntiSSEA1 1000/C+ | 10 | 9 | 9 | 4.9 |
| **TOTAL** | **50** | **42 (67-100%)** | **37 (56-90%)** | **4.7** |

*Ex-ovo dose-response study with deposition of antiSSEA1 and baby rabbit serum to the dorsal side of epiblast*. In all control embryos the PGCs counts were in the order of hundreds, while the percentage of embryos with scarce or without PGCs were increasing with 1/10 and 1/100 concentration of antiSSEA1. The dose-response study evaluated “in toto” revealed that antiSSEA1 100/C+ 1/40 was the most effective treatment destroying PGCs, because none of PGCs were seen in any embryo receiving this treatment (Table 2 and Fig 2c).

**Figure 2.**
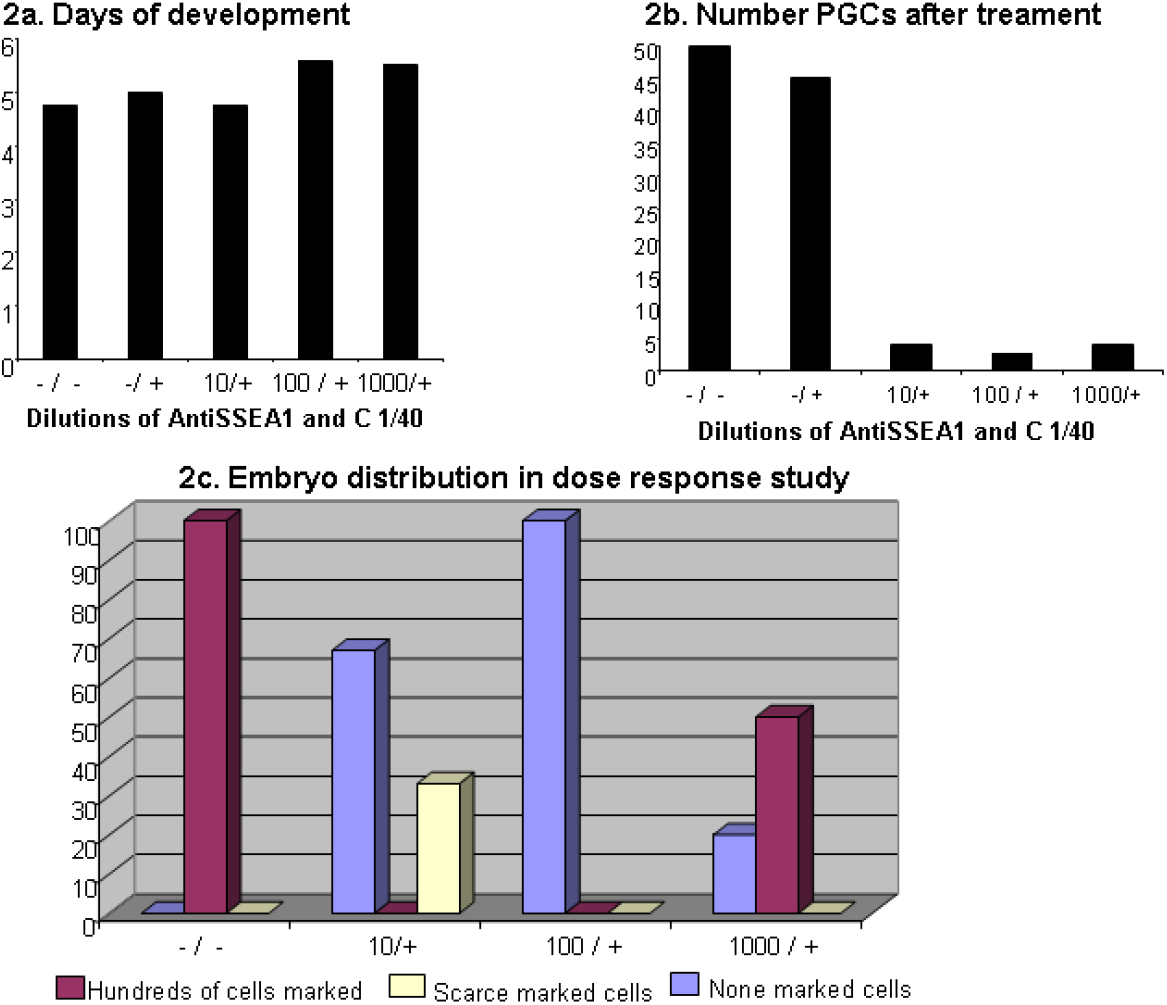
Treated embryos. The treatment was deposited to the dorsal side of epiblast. Days of survival (2a) and Number of PGCs (2b). PGCs were counted after a complete sectioning of embryos, a total of 2,400 tissue sections were evaluated. PGCs were immunochemically marked with antiSSEA1, biotinylated anti-mouse, followed by AP conjugated streptavidin and Fast Red. Three different experiments with at least 4 embryos per treatment group. 2c) Dose-response study in top-treated embryos, the PGCs were immunochemically marked with antiSSEA1 and antimouse alexa 488 “in toto”. Y-axis percentage of embryos in each treatment group. X-axis: AntiSSEA1 (-, 1/10, 1/100 and 1/100)/Baby rabbit serum or C (− or + at 1/40) designed as -/-, -/+, 10/+, 100/+ and 1000/+. Each group has at least three embryos.

**Table 2.**
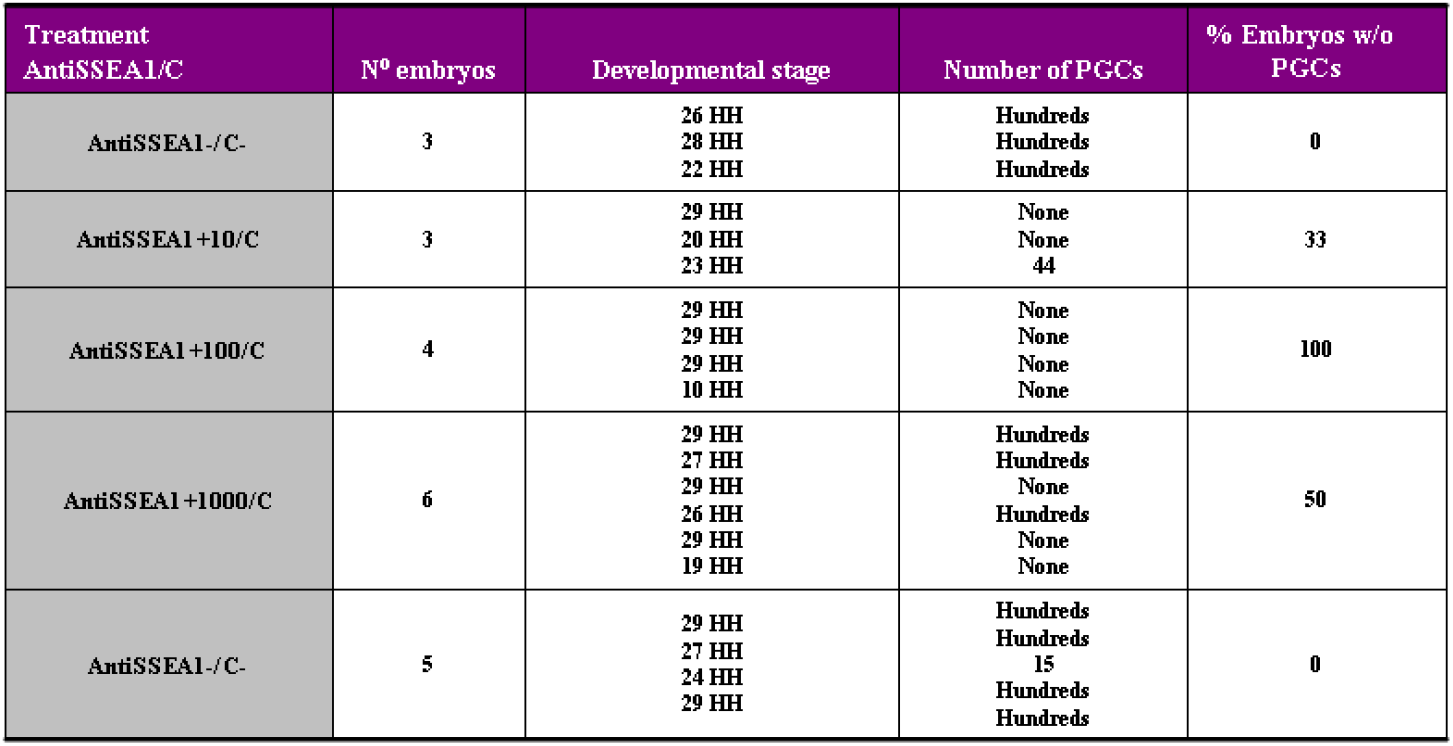
Results of dose-response study. The treatment was sequentially deposited to the dorsal side of epiblast. The majority of the embryos overpassed day 4 of development or 22 HH stage (Hamburger&Hamilton's classification). None embryo showed marked PGCs in the treated group antiSSEA1 100/C+.

*Embryos treated with injections into the subgerminal space*. The evaluation of embryonic survival revealed that the injected treatments and transfer manipulations did not affect neither embryo “early development” (67-100%) nor “late development” (50-75%) in any of the treated groups (AntiSSEA1 100/C+100, 100/C+1000 and /C+10000 groups) compared with controls (AntiSSEA1-/C- and AntiSSEA1-/C+1000), Table 3. The mean survival for injected embryos (4.4 days) was not different from top-dressed ones (4.7 days; Table 3 and Fig 3a). The number of PGCs detected was decreased in treated (41, 28 and 45 PGCs in AntiSSEA1 100/C+100, 100/C+1000 and 1000/C+10000 groups, respectively) compared with control embryos (82 and 100 PGCs in AntiSSEA1-/C- and AntiSSEA1-/C+1000, Fig 3b). The maximum cytolytic activity was achieved with anti SSEA1 100/C+1000. The complement alone did not show any cytolytic activity at 1/1000 dilution. Moreover, in a study of complement cytotoxicity all 6 embryos injected with AntiSSEA1-/C10 developed and had a high number of PGCs.

**Table 3.** Embryo distribution of “early” and “late development” after subgerminal injections in chicken embryos at stage X. “Early development” 24 hours after the application of the treatment, the embryo continues to advance. “Late development” at day fourth embryos with heartbeat and vascular tree well formed.

| Treatment<br>AntiSSEA1/complement | N° embryos | Early development | Late development | N° days |
| --- | --- | --- | --- | --- |
| AntiSSEA1-/C- CONTROL | 12 | 9 | 6 | 4.7 |
| AntiSSEA1+100/C+1000 | 10 | 8 | 8 | 4 |
| AntiSSEA1+100/C+100 | 7 | 6 | 4 | 3.7 |
| AntiSSEA1+100/C+1000 | 4 | 4 | 4 | 5 |
| AntiSSEA1+100/C+10000 | 6 | 5 | 5 | 4.7 |
| Cocktail | 6 | 3 | 0 |  |
| TOTAL | 43 | 35 (67-100%) | 27 (50-75%) | 4.4 |

Unexpectedly, when antibody and complement were injected together in a single injection, only half of the embryos reached “early development” and none “late development” (Table 3 and Fig 3a).

**Figure 3.**
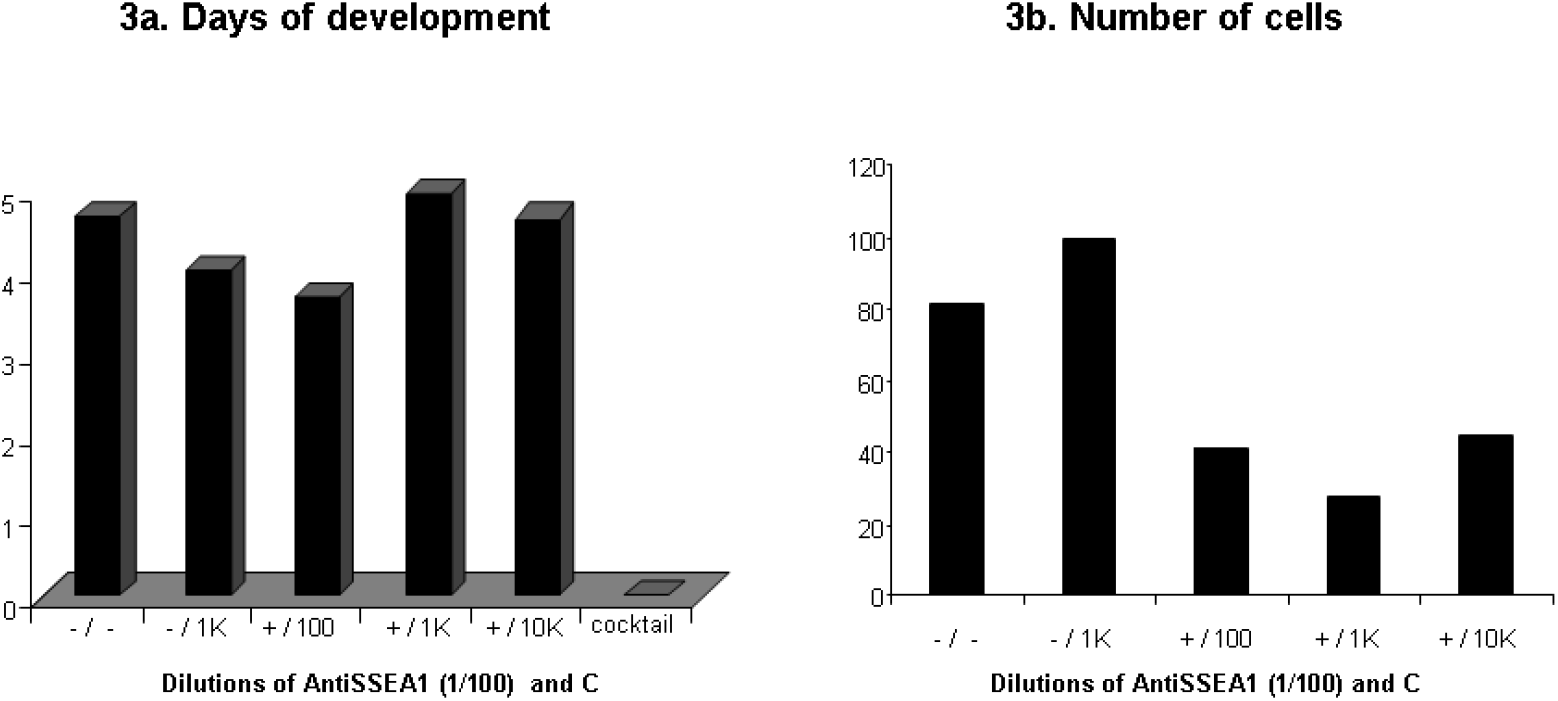
Subgerminally injected stage X embryos. Days of embryo survival (3a) and Number of PGCs (3b). The PGCs were counted after sectioning of entire embryos, a total of 3,900 tissue sections were immunostained and evaluated. 3a) Days of embryo survival after sequential injections of antiSSEA1 (- or + 1/100) and C (-, 1/100, 1/1K and 1/10K). Joint injection of antiSSEA1 and baby rabbit serum (cocktail), none of embryos were developed. 3b) Number of PGCs, the maximum cytolytic activity was archived between antiSSEA1 +/100-+/1K. The baby rabbit serum alone did not show any cytolytic action (-/1K).

### Immunocytochemistry and histological studies

*Immunohistochemistry in tissue sections*. In top-dressed and injected embryos, a total of 6,900 tissue sections were immunostained and the number of PGCs counted (Fig 4).

**Figure 4.**
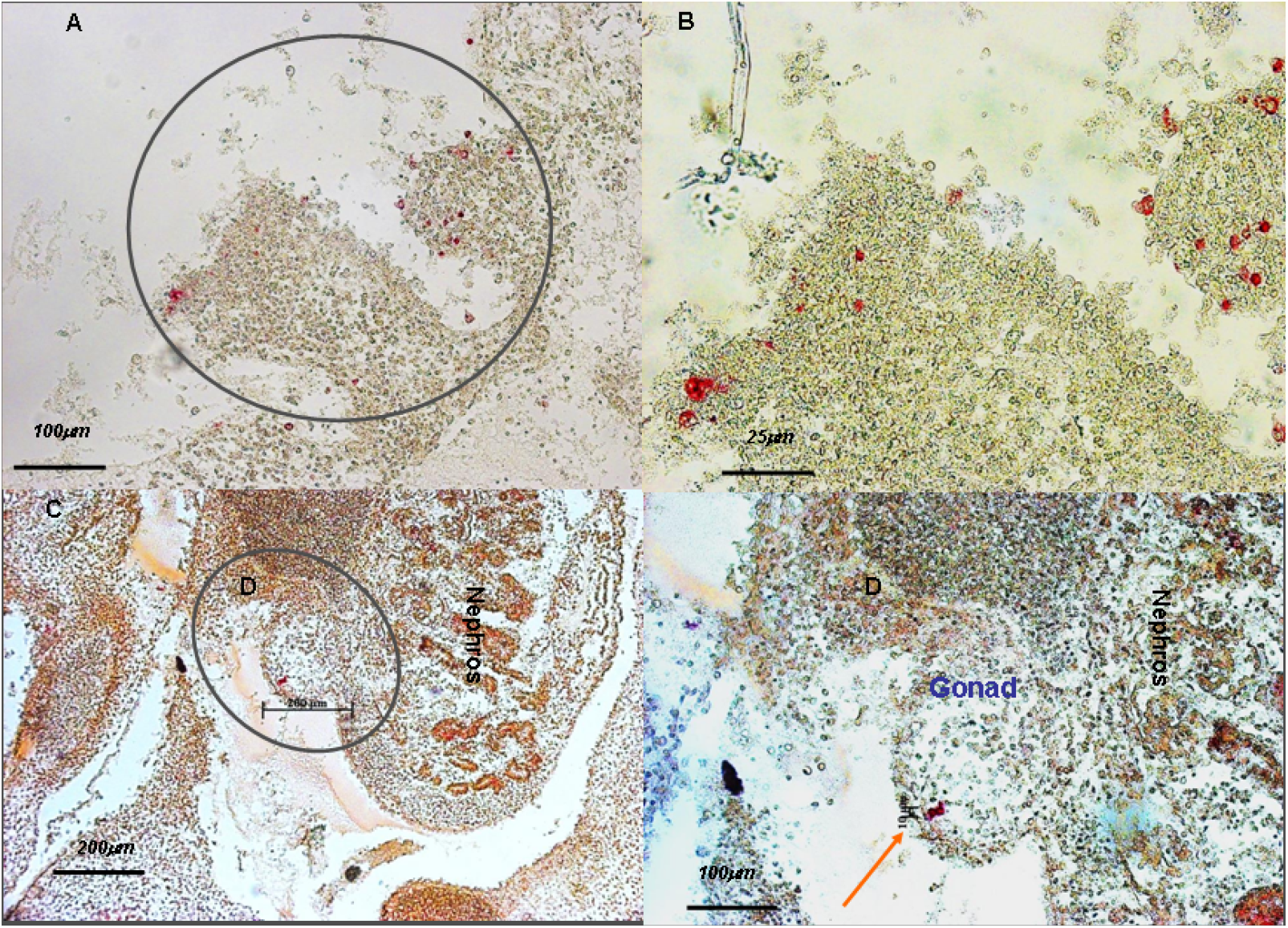
Histology of embryos treated with antiSSEA1 and baby rabbit serum. Top-dressed stage X embryos. Sections of gonads from control and treated embryos (5-days). A and B Control embryo PBS treated, only PGCs are immunochemically marked with antiSSEA1, biotinylated anti-mouse, followed by AP conjugated streptavidin and Fast Red (AX200 and BX400). C and D treated embryo showing one red cell colony (orange arrow), very few PGCs were counted in all sections. A total of 2,400 tissue sections were immunostained and analyzed. Haematoxilin counterstaining (C X100 and D X200).

The data “number of PGCs” have been already presented above (Fig 2b and 3b).

*Immunohistochemistry “in toto”*. In order to stain PGCs the embryos needed a strong permeabilization step (1% DMSO+0.1% triton X-100 in PBS overnight) and long periods of antibodies incubations (antiSSEA1 and Ig G donkey antimouse alexa 488, with 48 and 24 hours at room temperature, respectively). In control embryos the presence of hundreds of PGCs can be visualized in Fig 5 A. In Fig 5B and 5C most of the PGCs are already in the gonads but some are in their way migrating from the aorta, only PGCs were marked.

In some embryos of the antiSSEA1-/C+ group the immunocytohistology showed hundreds of labelled cells in their gonads but the labeling was not as strong as in the antiSSEA1-/C- group (Fig 5D E). The confirmation of the PGCs presence was made with a histological Alcian Blue-PAS staining (Fig 5F). In contrast, none of the four treated embryos showed any PGCs (Fig 6 A′, B′, C′ and D′).

**Figure 5.**
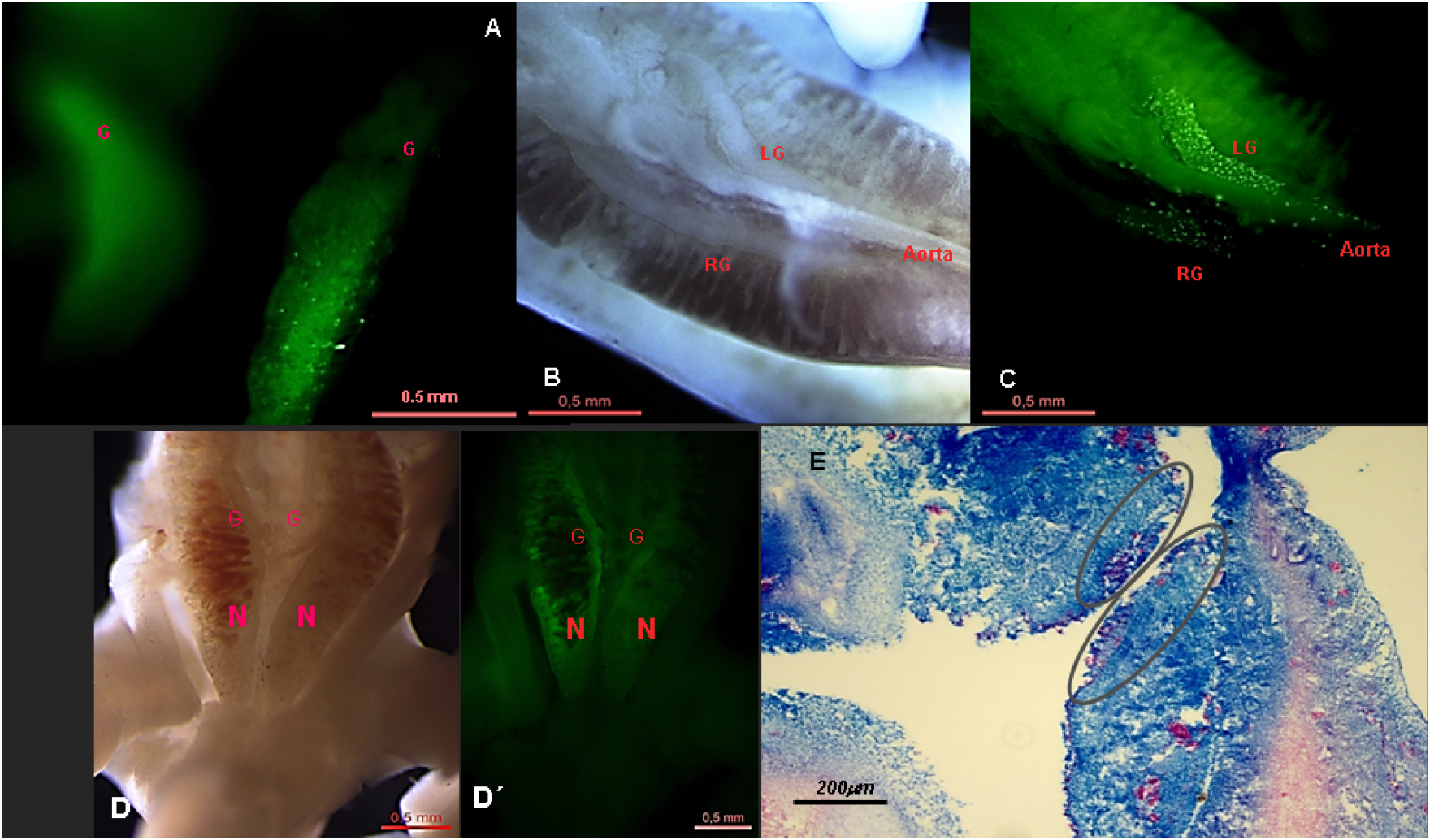
In vivo dose-response study in stage X embryos top-treated. The presence of PGCs was assessed by immunohistochemistry “in toto”. A control embryo only PGCs were stained. B and C, most of the PGCs are already in the gonads but some are in their way migrating from the aorta (C). D and D′ Nephros and gonads from an embryo treated with BRS (anti-SSEA1 -/C+). Leica MZIII. Embryos needed a permeabilization step (1% DMSO+0.1% triton X-100 in PBS overnight) and long periods of antibodies incubations (antiSSEA1 and Ig G donkey antimouse alexa 488, with 48 and 24 hours at room temperature, respectively). LG=left gonad, RG=right gonad and A=aorta. E) Histology of D embryo's gonads (grey circles), PGCs in the gonads are purple (PAS and Alcian blue positives). Embryo was Alcian Blue and PAS stained “in toto” and then sectioned (X100).

**Figure 6.**
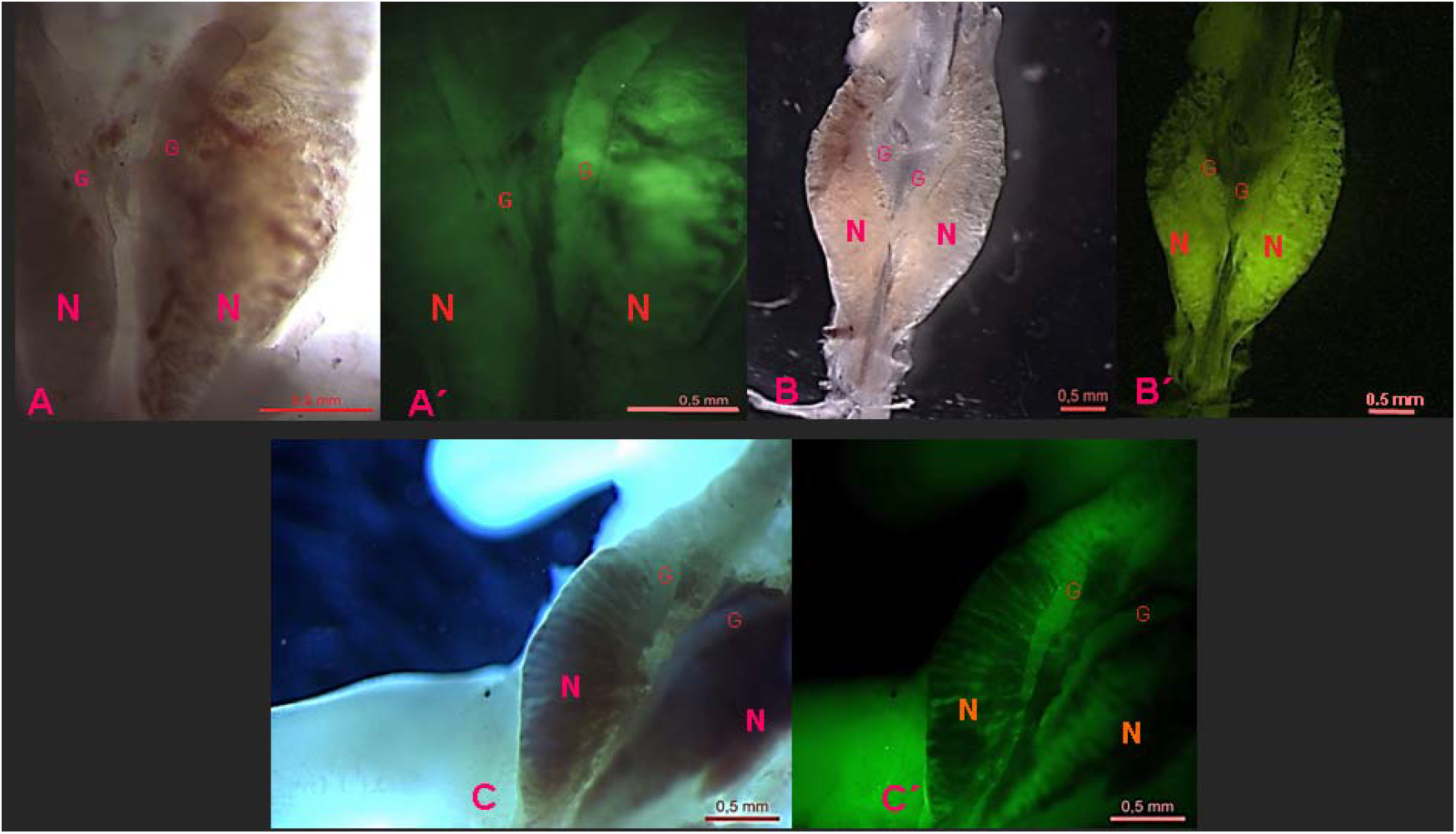
Treated embryos from “in vivo” dose-response study in stage X embryos top-treated. Embryo development did not progress beyond 6 days (HH 29). A B and C treated embryos with absence of PGCs. A B C and A′, B′, C′ visible light and fluorescence. AntiSSEA1 immunocytochemistry was made “in toto” with antimouse alexa 488. Embryos evaluated under fluorescence stereoscope Leica MZIII. G=gonad N=nephros.

## DISCUSSION

Our results demonstrated that immune-mediated ablation is an effective method for decimating the internal population of PGCs when preparing recipient chick embryos for germ line chimera construction. Germ cell numbers can be substantially reduced while embryo development is not affected. Treatment, dose and route had an impact on effectiveness and on survivability.

“In vivo” top-dress application, directly on the germinal disc, is, by far, the best assayed. No instrumentation is needed (stereomicroscope or micro injector) and it is the most gentle on the embryo. The response, in terms of PGCs numbers in the gonads, was dose dependent. Almost a complete ablation was achieved using antiSSEA1 and complement at 1/100 and 1/40 dilutions, respectively. These doses are coincident with our results obtained in the “in vitro” competition studies with SBRCs. We must emphasize that the duration of embryonic development and the organic structures were normal. There was a concern that this form of administration could be insufficient, as the molecules of antiSSEA1 and complement must cross not only the vitelline membranes but also the blastoderm itself. Gerhart et al. were able to ablate two lineages of epiblast cells expressing skeletal muscle (MyoD) and neuronal (NeuroM) surface antigens (Gerhart et al. 2008; Gerhart et al. 2010). Supporting in part, our studies in using this route, since PGCs are concentrated on the hypoblast at stage X.

This was the reason why direct treatment injection on the subgerminal space, that is, immediately below the hypoblast, was also assayed. We expected this route would be more effective on PGCs but also more damaging on the embryo because of repeated puncture of the vitelline membrane. As anticipated, relative to top-dressing, sequential subgerminal application required less complement (antiSSEA1/100 plus C 1/1000), to yield an equally extensive PGCs ablation; survival of the embryos was, however, not impaired. There could be two reasons why less complement is needed when injected; first, the effective complement concentration around the target PGCs should be undoubtedly higher, as losses due to diffusion are minor; second, the antiSSEA1 labelling on the target cells is more extensive, since the injected antibody is not washed away after a short reaction time as when top-dressed.

In both strategies, the natural Ig and innate complement system present in BRS could be implicated (Gerencer et al. 1998). The most probable mechanism would be that once antiSSEA1 is bound to the outer Galβ (1-4), Fucα (1–3)-GlcNAcβ1-R4 of PGCs, the rabbit innate complement system will consequently induce PGCs cytolysis by Complement Classical Pathway activation (C_1q_). Gerhart's results and the fact that in stage X embryos, neither complement enzymes nor IgM are present, support this mechanism. AntiSSEA1 has been frequently used to enrich PGCs isolates (Etches 1998 and Mozdziak 2006) although other antibodies have also been successful, as EMA-1, NC-1 and VASA. The later could be an optimum choice because of its specificity for germ line cells, as long as a good chicken antiVASA would be available.

We assayed a third administration strategy, joint injection of antiSSEA1 plus BRS into the subgerminal space, in case the double injection proved too detrimental to embryo viability. An unforeseen and remarkable disruption was observed on germinal disc development following injection of the antiSSEA1-BRS mixture. It suggests that a massive, untargeted cytolytic activity was developed that damaged several embryonic cell populations and structures. The only report on simultaneous use of antibody and serum (premixed before administration) “in vivo” is that of Takami et al. (2006) when treating a CNS lymphoma with rituximab, after previous treatment failures applying the antibody alone intraventricularly (Schulz et al. 2004). Generally, when the antibody-complement complex is used to direct a cytolysis over certain cell populations, the “modus operandi” is: the cells are first exposed to the antibody and, later on, the complement is applied to induce the lysis of only those cells previously targeted (Gerhart et al. 2008, Gerhart et al. 2010, Chen and Melton 2007). Maybe other enzymes besides Cq1 of the complement system could be activated through the Lectin pathway. Several types of lectins present in serum have been studied, among them, mannose-binding lectins (bind to mannose) and ficolins (bind to GlcNAc or fucose). The latter need IgM to activate Lectin pathway (Endo et al. 2015). Humoral ficolins present in BRS could very well bind to IgM (antiSSEA1) when mixed and subsequently activate the complement enzymes (C_4_ and C_2_), which in turn, will destroy indiscriminately blastodermal cells. Supporting our hypotesis, Lei et al. (2015) have described a novel IgM–H-Ficolin complement pathway which ablates “in vitro” allogenic human cancer cells. Glycans covering blastodermal cells, as well as cancer cells, have GlcNAc and fucose (Endo et al. 2015 and Lei et al. 2015). In stage X chick embryos the glycocalyx of blastodermal cells and PGCs are richer in GlcNAc than in mannose (author unpublished observations). Taken together all we suggest a possible implication of IgM-ficolins complexes in this massive destruction. However, obviously the question of why antibody-complement together strongly increased cytolitic efects needs to be further investigated.

The use of monoclonal antibodies and complement is one of the most promising therapies in cancer. However, in long term treatments with specific antibodies, some tumours produce their own complement and others modulate the complement regulators resulting in the inhibition of the innate complement. As far as we know, premixing of exogenous complement and monoclonal antibody has not been reported as a strategy to boost antibody therapeutical effects. Our results are supported by the reported treatment of primary central nervous system lymphoma in humans using Rituximab (monoclonal antibody against primary lymphoma cells) and exogenous complement (Nishimura et al. 2003). Therefore, the mixture of antibody-complement previously to its local application could be very useful to treat large established tumours from which available antibodies exist and also to potentiate the antibody efficiency.

On the other hand, the lack of cytolytic response in our control, demonstrates that complement alone, even when directly injected into the subgerminal space at the highest concentration (C+1/10 dilution), is not cytotoxic for the developing stage X embryo: it failed to reduce the endogenous PGCs population and it did not affect embryo viability at all.

Therefore, two protocols successfully produced lysis of the primordial germ cells by specific cell labelling with antibody followed by heterologous serum with innate complement system present in BRS. Dress-on application is simpler and faster than subgerminal injection while equally effective, although larger quantities of reagents are needed. Both methods are capable of producing an almost complete ablation of the germline cells in chicken embryos with a high efficiency without damaging other embryonic structures at stage X embryos.

## MATERIALS and METHODS

### Egg source

Freshly fertilized laid chicken eggs were obtained from Cobb SA (Alcalá de Henares, Madrid, Spain). Housing and management of the laying hens (Gallus gallus, Linn) comply with welfare and sanitary EU standards.

### “In vitro” cytolysis of avian blastoderm cells

*Competition tests of dispersed blastodermal cells with haemolytic system (HS)*. The optimal dilution of the available complement systems was established in a complement assay with baby rabbit serum (BRS) and haemolytic system (HS; SRBC and haemolysin in gelatine-Veronal buffer containing Ca^2+^ and Mg^2+^, SRBCs). All reactives were generously supplied by Margarita López Trascasa, Hospital La Paz, Madrid Spain. Indirect estimation of blastoderm cytolysis was made using SRBCs, HS and blastoderm cells in competition assays. Maximum haemolysis was induced by addition of distilled water to HS. The CH_50_ was established as the range of those dilutions of BRS when haemolytic action was exhausted, and SRBCs started to settle down in round-bottom ELISA plates.

A total of 32 blastoderms were isolated from embryonated eggs at oviposition and disaggregated with repeated pipetting for three competitive studies. First, the blastoderm cell suspensions of ten stage X embryos (300 μl approx 10^7^ cells/ml) were incubated in assay tubes with monoclonal antibody antiSSEA1 obtained as concentrate from the Developmental Studies Hybridoma Bank (1/100 and 1/1000) during 45 minutes at 37°C in a water bath. Next, BRS (1/10; 1/100; 1/1000 and 1/10000) plus HS were added and, after incubation for an additional hour at 37°C in a water bath, the reaction cocktail was centrifuged 1000 g for 10 minutes and the supernatant was transferred to a multi-well plate where absorbance at 405 nm was recorded. At the end of the second incubation, each reaction tube was evaluated for SRBC sedimentation: a reddish sediment in the tube reveals incomplete haemolysis since the complement in the reaction mix has been exhausted. Conversely, absence of SRBC sediment denotes excess complement activity in the mix. The degree of haemolysis was measured as the absorbance of the supernatants at 405nm. Saturation curves revealed that CH_50_ was reached after adding BRS between a range of 1/10 −1/100 dilutions.

Next, a total of 26 entire blastoderms were isolated for the evaluation of cell viability described previously (Gerhart et al. 2008). In brief, each isolated embryo was placed in a well with cell culture medium M199 in four experiments following previously described culture conditions (Lopez-Diaz et al. 2016). Next, the blastoderms were challenged to antiSSEA1 (1/100 and 1/1000) incubated during 45 minutes at 37°C and then to BRS (1/50). After incubation for an additional hour at 37°C, the evaluation of cell destruction was made adding 4% trypan blue (4μl). Control blastoderms were incubated with only M199 or BRS+M199.

### Ex-ovo treatments applied to stage X embryos and incubated in surrogated egg shells following Perry’s system II

*Deposition of antiSSEA1 and baby rabbit serum to the dorsal side of epiblast*. Embryonated chicken eggs at ovoposition (stage X) were cleaned with alcohol or with a solution of sodium hypochlorite for 30 or 60 minutes before use. The eggs content was poured on sterile Petri dishes to expose and manipulate the embryos following Gerhart’s procedure (Gerhart et al. 2008) with some variations. Briefly, the albumen covering the germinal disc was cleaned with sterile cellulose tissues until a well-marked hollowness appeared. Once the germinal discs were free from albumen, 100 μl of antiSSEA1 (1/10, 1/100 and 1/1000) were pipetted on top of the germinal disc and covered with a piece of parafilm. The treated embryos were incubated in an oven at 37°C saturated with moisture for 1 hour. The excess of unbound monoclonal antibody was washed away three times with PBS and BRS was immediately top-dressed on the blastoderm. Baby rabbit serum (diluted 1/40 in PBS + 0.1% skim milk) was used as a source of complement because it only has the innate complement system (C) and natural immunoglobulins, lacking immune IgG and IgM (Gerencer et al. 1998). The blastoderm was covered with parafilm and incubated for another hour in the same conditions. After washing the BRS, the contents of eggs were transferred to surrogate egg shells following Perry’s system II (Perry, 1988). A total of 50 embryos were evaluated and histologically studied from three different experiments with at least 4 embryos per treatment group (antiSSEA1 -/C-; antiSSEA1 10/C+, antiSSEA1 100/C+, antiSSEA1 1000/C+ and antiSSEA1 -/C+).

*Treatment with two sequential injections into the subgerminal space*. Embryos were injected into the subgerminal space where PGCs are localized at stage X. First, a volume between 5-10 μl of antiSSEA1 (1/100) was injected with micropipettes using a pneumatic fluid microinjector (Injectmatic, Geneva, CH). The treated embryos were incubated in an oven at 37 °C saturated with moisture for 1 hour. Later, BSR was injected at different dilutions (1/100, 1/1000 and 1/10000). After the last injection the contents of eggs were transferred to surrogate egg shells. A total of 43 embryos were evaluated from three different experiments with at least 4 embryos per treatment group (antiSSEA1 -/C-, antiSSEA1 100/C100, antiSSEA1 100/C+1000, antiSSEA1 100/C+10000 and antiSSEA1 -/C+1000). In order to rule out a possible direct cytotoxic effect of BRS, a study was undertaken injecting subgerminally a maximum dose of complement 1/10 (antiSSEA1 -/C+10), including as well, a control group (antiSSEA1-/C-).

*Treatment with a single injection into the subgerminal space*. Six embryos were injected into the subgerminal space as indicated above, except that antiSSEA1 and BRS were jointly applied in a single injection (antiSSEA1 10/C+10).

*Evaluation of embryo development*. Everyday “embryo development” was checked and recorded. Embryo development was recorded as “early development” on progress, if next day after the application of the treatment, germinal disc showed normal growth and no yolk leaking was seen. On third day “late development” was recorded as normal when the vessels were formed and the heart was beating. Embryos were incubated at least until 5-6 day in an egg incubator (Octagon) on system II (Perry 1988). Once the incubation was over or the embryo development stopped, the embryos were dissected, fixed and conserved in methanol 70% until histological and immunohistological studies were undertaken.

### Immunocytochemistry and histological studies

*Top-dressed and sequential injections into the subgerminal space embryos*. The presence of PGCs was assessed by an alkaline phosphatase (AP) immunocytochemical detection system using antiSSEA1 (DSHB; Iowa City, IA, U.S.A.). Briefly, embryos were fixed in Bouin’s solution overnight, embedded in parafine and entirely cut with microtome into tissue sections of 10 μm. A total of 6,300 tissue sections were analyzed (a mean of 100 tissue sections per embryo). To minimize nonspecific binding, the tissue slices were treated for 3 hours with 3% BSA and 0.1% triton X-100 in PBS before immunostaining. The optimal concentration of each antibody was selected based on the results of preliminary experiments (1/300 for antiSSEA1 and anti-mouse). Tissues were incubated for 24-48 hours with the primary antibody at 4°C and subsequently reacted for 24 hours each with biotinylated anti-mouse (Sigma B0529, for monoclonal first antibody) followed by AP conjugated streptavidin (Sigma E2636). Fast Red was used as substrate chromogen. To avoid interference by the potential endogenous activity of AP, the tissues were treated with 0.02 M levamisole or heat inactivated at 70 °C for 30 minutes.

*Dose-response study*. In the dose-response study the number of PGCs was evaluated “in toto” embryos by immunohistochemistry. Briefly, embryos were fixed in 4% paraformaldehyde in PBS overnight at 4 °C, permeabilized using 1% DMSO and 0.1% triton X-100 in PBS overnight and unspecific binding blocked at 4 °C overnight with 3% BSA and 0.1% triton X-100 in PBS before immunostaining. The presence of PGCs was assessed by Ig G donkey antimouse alexa 488 (24 hours at room temperature, Invitrogen) and antiSSEA1 as primary antibody (48 hours of incubation). This immunohistochemistry allowed us a clear identification of PGCs with absent fluorescent background. The nephros and gonads were dissected and the number of PGCs twice counted under a stereoscope LEICA MZIII and an inverted fluorescence microscope by two people (Nikon, Eclipse TE300; Tokyo, Japan, fitted with a H910104A, TE-FM Epi-Fluorescence Lamp). The results were classified as: no cells = none PGCs marked, countable cells = less than 100 marked PGCs and uncountable cells = hundreds of PGCs were marked. In order to confirm that PGCs were marked, other histological studies were made like PAS. In some embryos with weak fluorescece labelled PGCs, specially all those treated only with complement, the gonads were stained with Alcian Blue and PAS (Merk Germany 1.016460001) following the manufacturer’s instructions. PGCs are Alcian Blue and PAS positive which gives a unique purple color at HH 27 developmental stage (5 days).

## Acknowledgements

The authors acknowledge the valuable support of Dra Capitolina Díaz Martínez, President of AMIT (Asociación de Mujeres Investigadoras y Tecnólogas). The valuable help of Soraya Martínez-Alcocer and Diana González-Chamorro for histological processing is gratefully acknowledged. This study was partly supported by the grant AGL 2009E06345-MICINN (Spain).

The monoclonal antibody antiSSEA1 developed by Solter and Knowles was obtained from the Developmental Studies Hybridoma Bank developed under the auspices of the NICHD and maintained by The University of Iowa, Department of Biology, Iowa City, IA 52242.

## Role of the founding source

This study was supported by the grant AGL 2009E06345–MICINN (Spain).

